# Generative Adversarial Networks Can Create High Quality Artificial Prostate Cancer Magnetic Resonance Images

**DOI:** 10.1101/2022.06.16.496437

**Authors:** Isaac R. L. Xu, Derek J Van Booven, Sankalp Goberdhan, Adrian L. Breto, Mohammad Alhusseini, Ahmad Algohary, Radka Stoyanova, Sanoj Punnen, Anton Mahne, Himanshu Arora

## Abstract

**Purpose:** Recent integration of open-source data to machine learning models, especially in the medical field, has opened new doors to study disease progression and/or regression. However, the limitation of using medical data for machine learning approaches is the specificity of data to a particular medical condition. In this context, most recent technologies like generative adversarial networks (GAN) could be used to generate high quality synthetic data that preserves the clinical variability.

**Materials and Methods:** In this study, we used 139 T2-weighted prostate magnetic resonant images (MRI) from various sources as training data for Single Natural Image GAN (SinGAN), to make a generative model. A deep learning semantic segmentation pipeline trained the model to segment the prostate boundary on 2D MRI slices. Synthetic images with a high-level segmentation boundary of the prostate were filtered and used in the quality control assessment by participating scientists with varying degree of experience (more than 10 years, 1 year, or no experience) to work with MRI images.

**Results:** The most experienced participating group correctly identified conventional vs synthetic images with 67% accuracy, the group with 1 year of experience correctly identified the images with 58% accuracy, and group with no prior experience reached 50% accuracy. Nearly half (47%) of the synthetic images were mistakenly evaluated as conventional images. Interestingly, a blinded quality assessment by a board-certified radiologist to differentiate conventional and synthetic images was not significantly different in context of the mean quality of synthetic and conventional images.

**Conclusions:** This study shows promise that high quality synthetic images from MRI can be generated using GAN. Such an AI model may contribute significantly to various clinical applications which involves supervised machine learning approaches.

## Introduction

Prostate cancer is the most common cancer among men and second most common cause of cancer related death for men in the US^1^. Relatively high incidence of prostate cancer has invoked discussion on a national screening program ^2^. The National Cancer Institute alone was estimated in 2020 to have spent $209.4 million on prostate cancer research and $233.2 million on clinical trials ^3^. Despite this, the costs of treating prostate cancer are increasing more rapidly than those for any other cancers ^4^. Multiple studies have suggested that the high cost and time investment of prostate cancer clinical trials pose a significant barrier to research, as well as the formation of large cohorts of patients with multiple years of follow-up for high-value conclusions ^5^.

The use of Magnetic Resonance Imaging (MRI) in prostate cancer research and treatment is effective for prognosis, diagnosis, active surveillance, and reduced need for biopsy procedures in lower risk patients ^6,^. Additional benefits of MRI include ease of visualization, staging, tumor localization, risk stratification, active surveillance monitoring, and detection of local failure after radiation therapy ^7^. Prostate MRIs provide more clear and detailed images of soft-tissue structures of the prostate gland than other imaging methods, making them valuable data for research ^,,8^.

Ongoing advancements in machine learning technologies (or Artificial Intelligence, AI) have contributed to scientific discovery in medical imaging and show promise for its future uses ^9^. For example, radiologists typically need many years of practice to read MRI scans accurately, leading to disagreements between radiologists looking at the same scan. However, AI has helped make the learning curve easier for practicing radiologists and lowers the error rate by performing tasks such as predicting aggressive cancer, or AI segmentation of organs. When applied to diagnostic imaging, AI has shown high accuracy in prostate lesion detection and prediction of patient outcomes, such as survival and treatment response ^10^. Machine learning can help the medical experts in early disease detection; for prostate cancer, for example, machine learning classifiers predicted Gleason pattern 4 with approximate if not less accuracy than experienced radiologists ^11^. Other deep learning approaches such as convolutional neural networks (CNNs) and generative adversarial networks (GANs) have been used in various applications of medical image synthesis of PET, CT, MRI, ultrasound, and X-ray imaging, particularly in the brain, abdomen, and chest ^12^. A recent study using synthetic MRI in colorectal cancer also found no difference in signal-to-noise ratio, contrast-to-noise ratio, and overall image quality between conventional and synthetic T2-weighted images ^13^. GANs have also been shown to be able to create realistic synthetic brain MRIs ^14^. All these facts combined contribute to our hypothesis that we can use deep learning image synthesis to create medical images that translate into clinical use and avoid huge costs associated with repeat follow-ups. Synthetic prostate MRI using GANs may be a future possibility for large scale machine learning projects, where millions of image samples can be generated without the need for millions of patients.

## Materials and Methods

### Prostate MRI Images

Prostate MRI images used in this study were obtained from three sources: (1) T2W tse (T2 weighted Turbo-Spin-Echo) images were downloaded from the first 60 patients of the Cancer Imaging Archive’s ProstateX challenge repository ^15^. (2) T2W fse (T2 weighted Fast-Spin-Echo Transversal) images including manually segmented contour data from 45 patients enrolled in fusion database, University of Miami’s Sylvester Cancer Center. (3) T2W fse images from 79 patients, enrolled in fusion database, University of Miami Department of Urology. All images used were displayed in the axial plane. All human investigations were carried out after the IRB approval by a local Human Investigations Committee and in accord with an assurance filed with and approved by the Department of Health and Human Services. Data has been anonymized to protect the privacy of the participants. Investigators obtained informed consent from each participant.

### Image Preprocessing

Digital Imaging and Communications in Medical Science (Dicom) information for prostate MRIs from each dataset was converted into nearly raw raster data (NRRD) format for better data processing, normalized to intensity values from 0-1, and then saved into 2D images from slices of the prostate along the z-axis using Python’s Matplotlib ^16^package. To control for MRIs of different dimensions and pixel sizing, images were cropped proportionally to the smallest dimension (x or y) into a square and then resized to a 500 × 500-pixel image using Python’s Pillow package.

### Synthetic Image Generation

For each prostate, the 2D image used to train a generative model was the slice from the z-axis index with the largest manually identified prostate. Synthetic images were created using Generative Adversarial Networks for each preprocessed image with SinGAN ^17^ [https://github.com/tamarott/SinGAN] under default settings. Using the 500 × 500 images in SinGAN would result in ten generative models for each image. The random sample synthetic image that was produced in the 8^th^, 9^th^, and 10^th^ edition of the models was considered for further analysis. SinGAN was used because of its ability to create a generative model off a <u>single</u> image, allowing for an unsupervised model, independent of the number of training images. For this study, 139 prostate MR images were trained using SinGAN to make unique generative models, with three synthetic samples being taken from each model.

SinGAN created synthetic images that maintained resolution of conventional MRIs. The images created used generative adversarial networks, invented by Goodfellow et al ^18^. SinGAN takes advantage of two neural networks, a Generator to generate image samples and a Discriminator to discriminate between conventional and synthetic samples. The training was performed on one image in a course-to-fine manner to make a model that can generate random samples (Figure 1). Synthetic images appear similar to the training image topologically, but have noticeable realistic variation, especially in the prostate and peripheral zone (PZ). Prostate and peripheral zone image and radiomic statistics are commonly used in studies involving prostate cancer risk, progression, tumor analysis, and clinical trial studies using radiation therapies ^19^. This is due to prostate cancer appearing as hypo-intense signal compared to higher signal depicting normal prostatic tissue ^20^.

**Figure 1:**
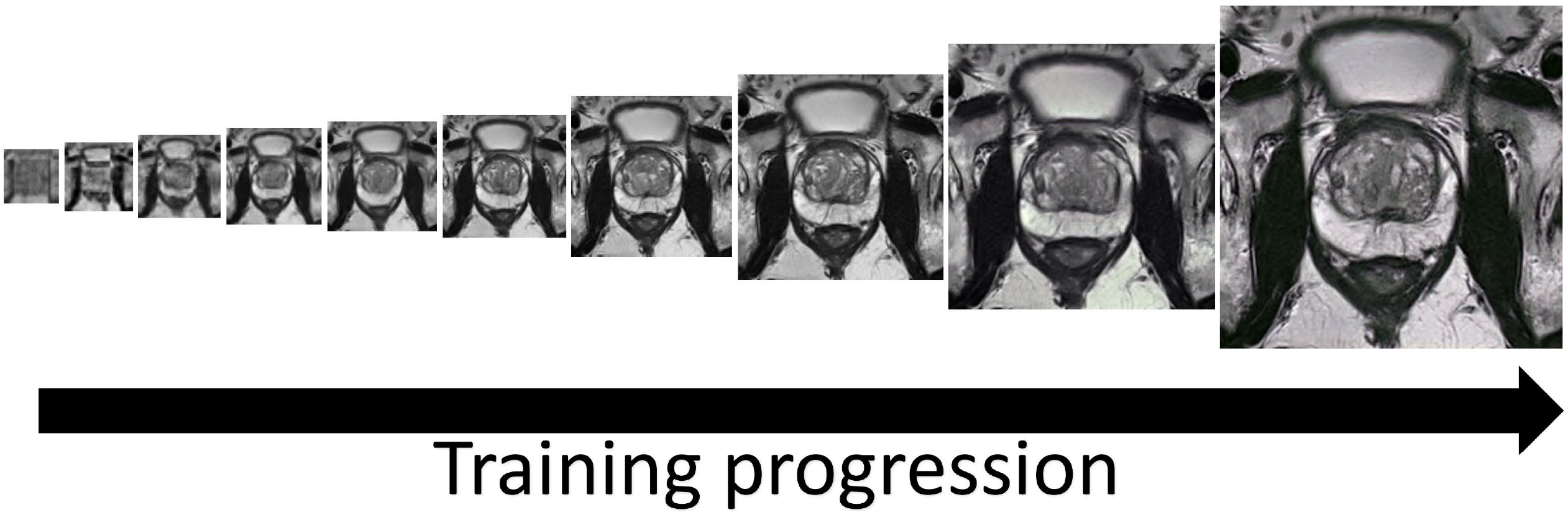
Image generation using SinGAN. SinGAN trains a model using generative adversarial networks in a course-to-fine manner. At each scale, the generator makes sample images that cannot be distinguished in down-sampled training images by the discriminator. The 8^th^, 9^th^, and 10^th^ scale images (the three rightmost images) of the model were resized and used in all quality control tests.

### Deep Learning Image Segmentation

Deep learning image segmentation of prostate MRI has been used in the past years with relative success ^21^. We hypothesized that high quality synthetic images can be successfully segmented by a neural network. Therefore, an image segmentation neural network was trained on the prostate contours from the dataset of 45 patients from University of Miami’s Sylvester Cancer Center. For each patient’s prostate MRI, the three middle slices of the contour and corresponding T2 image were used to train the network, totaling 135 images in the training set. The neural network was trained using the PyTorch ^22^ machine learning package and the code was adapted from existing code ^23^. The learning rate for each step of the model’s gradient descent was 0.00001 and the batch size was 3.

The outputted synthetic images were subjected to a quality control check using the trained model. For the synthetic image to pass the deep learning image segmentation quality check, the output prediction boundary for the prostate had to have been at least 10,000 pixels and be a single, unbroken boundary. The rationale for 10,000 pixels comes from our training dataset having a minimum contour pixel size of 10932 pixels.

### Quality Control Study

To test the realism of the synthetic images, a blinded quality control test was given to selected participants with varying expertise in prostate cancer MRI research. The first group was 2 scientists considered experts in the field with 9 and 13 years of experience with prostate cancer MRIs. The second group consisted of 2 scientists with approximately 1 year research experience in prostate cancer MRIs, which we classified as having some experience but not yet expertise. A negative control was given to a third group consisting of 4 scientists with no experience working with prostate cancer MRIs. The first quality control test was used on synthetic images that were manually selected from the output of the trained models from the 79 images obtained from the Urology department training set. In this first test, high quality synthetic images were manually selected, while synthetic images with high distortion were excluded. A total 60 images were used for this test, including 25 conventional images and 35 synthetic images. The radiation oncology team members were given five seconds to study the image, then recorded their prediction as conventional or synthetic.

To improve the fidelity of the study, a second quality control test was given to the same participants to test for variance, and to see if with more exposure they would have learned to differentiate better conventional and synthetic images. The images used this time were subjected to the deep learning image segmentation pipeline and then randomly selected without replacement for inclusion to both automate the process and control for human bias in image selection. Synthetic images that were generated from images <u>**not**</u> in the training data and passed the neural network criterion were pooled and randomly selected for this test. The test consisted of 26 conventional images and 36 synthetic images, and participants were given the same format as the first assessment.

## Results

### Deep Learning Image Segmentation

With a goal to create a pipeline that automates a process for determining if a synthetic image was high enough quality to be included for grading by fellow scientists and in future projects, as well as to test whether synthetic prostate MRIs can be used in similar machine learning contexts as conventional prostate MRIs, the image segmentation neural network model was trained on 135 images from 45 patients for 3000 steps. For each step of the training, a T2W image and corresponding contour were given as inputs (Figure 2A). The contour is a binary file of the same dimensions as the MRI. After 100 epochs of training, the neural network predicted the contour with a Dice similarity coefficient 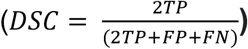 of 0.8857 and accuracy 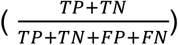 of 0.9795 (Figure 2B). The 1000^th^ step predicted with DSC of 0.9574 and accuracy of 0.9945. The 3000^th^ step predicted with a DSC of 0.9991 and accuracy of 0.9988. Although the model over-fit to the training data, we still saw an increase in the accuracy of the prediction on synthetic images with more epochs of training and decided to use the 3000^th^ step model for this analysis. For a synthetic prostate MRI image to pass the neural network segmentation pipeline’s criteria, the predicted segmentation had to have 10,000 pixels (4%) and be one unbroken contour (Figure 2C). The 10,000-pixel cutoff was based off the training data contours minimum 10932 pixels (4.3%) and median 19481 pixels (7.8%). The segmentation pipeline performed well in not predicting segmentation in synthetic images which subjectively had obscured and malformed prostates. Out of 654 synthetically generated images, close to 39% (253, or 38.6%) passed the neural network pipeline’s criteria. The images that did pass the criteria were used in the further analysis of this study (Figure 2D).

**Figure 2:**
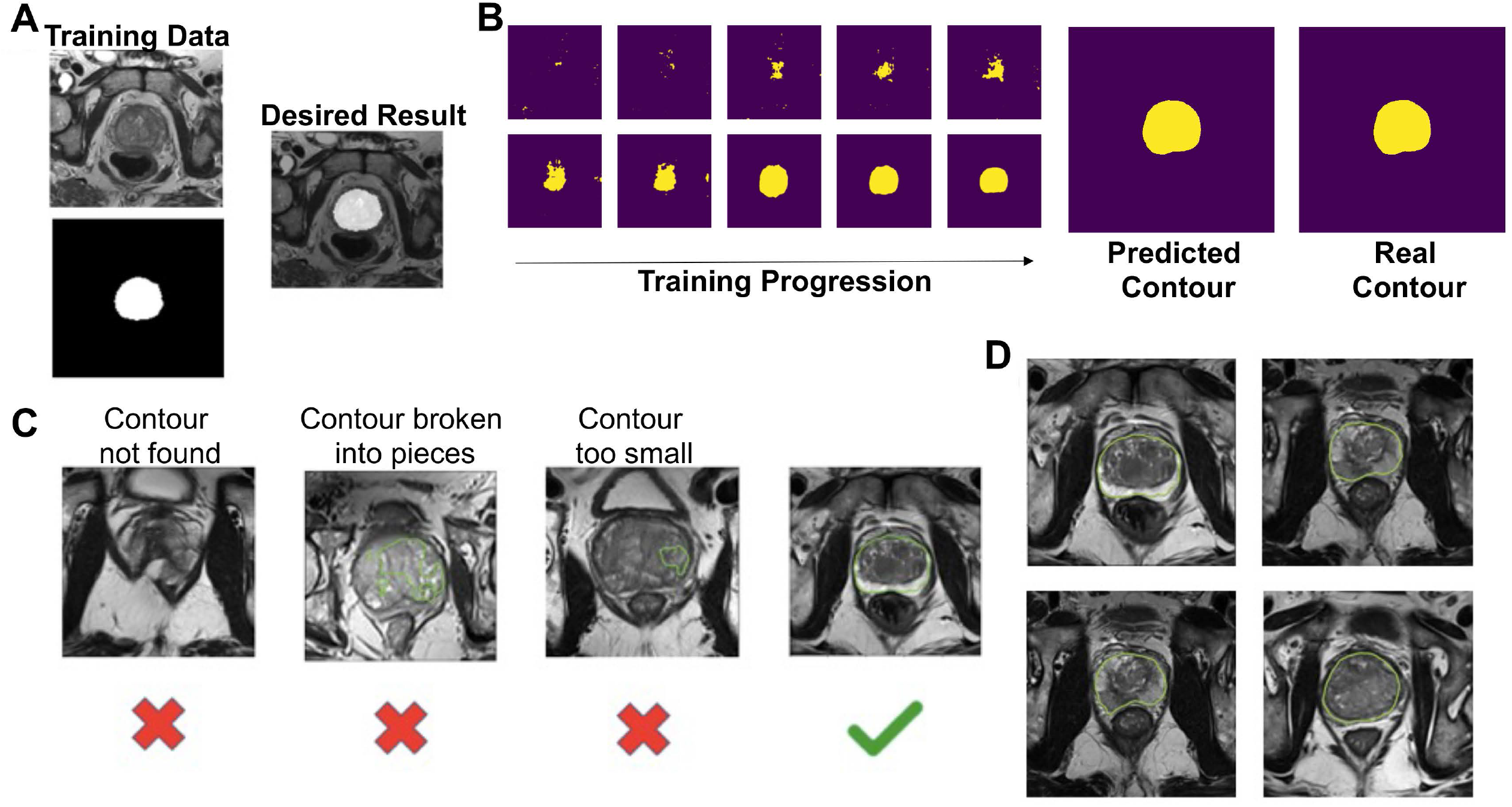
A) Example of training data for the image segmentation neural network and the desired result. Top left is the T2 tse Tra training image and bottom left is the corresponding binary contour. The deep learning image segmentation training progression shown for one image of the training set. The final predicted contour had a dice similarity coefficient of 0.99 to the real contour. To pass the deep learning segmentation pipeline, the predicted contour had to be one continuous contour greater than 10,000 pixels. D) Examples of the 253 synthetic images that passed the deep learning segmentation pipeline, shown with the prediction boundary.

### Round 1 Quality Control Test

The first-round quality control test was given to selected participants based on their experience with prostate cancer MRIs. Two participants had roughly a decade of experience (9 years, 13 years), two participants had about a year of experience, and four participants had no experience. Participants were shown 60 hand-picked images both synthetic and conventional and asked after 5 seconds to give their opinion if the image was synthetic or conventional. Results from this experiment are summarized in Table 1. The accuracy for groups were 62%, 55%, and 53%, respectively. A Pearson’s Chi-square test showed no significant association between experience level and the number of correct predictions (p=0.2855), no association between experience level and the number of false negatives (p=0.4125), and no association between experience level and number of false positives (p=0.6675); however, a significant association was found for concordance within groups when all groups were considered. No significant association in concordance was determined when comparing the participants with 10 years of experience to participants with 1 year of experience. These results demonstrate that for all levels, correctly identifying the difference between synthetic and conventional prostate MRI images is challenging.

**Table 1:** The results from the first and second round of quality control tests. The second round used the deep learning segmentation pipeline and random sampling for the test creation. False positives are defined as conventional images that participants labeled as synthetic. False negatives are defined as synthetic images which participants labeled as conventional. Concordance between the participants is defined as the proportion of answers that were the same between them.

|  | Round 1 |  |  | Round 2 |  |  |
| --- | --- | --- | --- | --- | --- | --- |
| **Amount Conventional** | 25 |  |  | 26 |  |  |
| **Amount Synthetic** | 35 |  |  | 34 |  |  |
| **Experience Level** | 10 Years | 1 Year | No Experience | 10 Years | 1 Year | No Experience |
| **% Correct** | 62 | 55 | 53 | 67 | 58 | 50 |
| **% FP** | 46 | 46 | 54 | 25 | 35 | 47 |
| **% FN** | 33 | 44 | 42 | 40 | 47 | 54 |
| **% Concordance** | 67 | 57 | 42 | 80 | 60 | 30 |

### Round 2 Quality Control Test

Figure 3 shows the workflow of this second quality control test. The second quality control test was to eliminate human bias in image selection (selecting the most realistic synthetic images and the lower quality conventional images), to automate the process, and to test if the participants learned the difference since participating in the first test. The second quality control test was given to the same team of scientists. Out of 654 generated synthetic images, 253 passed the criteria from the deep learning image segmentation pipeline and were randomly selected for the test. Table 1 illustrates the results of our quality control tests. Our three groups of participants scored correctly 67%, 58%, and 50%, respectively. A Wilcoxon signed ranked test shows the amount correct between participants’ first and second survey is not statistically different (p=0.725), regardless of group. A Pearson’s Chi-Square test shows that in there is an association between experience and group when considering all three groups (p=0.005682) however, there was no significant association between those with 10 years of experience and those with 1 year of experience (p=0.1824). Furthermore, the group with no experience studying prostate cancer MRIs had high variance between scores, with the highest score matching the average of the expert group (67%), yet the lowest score was 35%. There was no significant association between false negatives (p=0.2766) or false positives between experience levels (p=0.06051). These results mirror a similar study where expert and basic prostate radiologists found no difference in diagnostic performance between conventional MRI and synthetic MRI ^24^. They also show a similar result to another study where brain synthetic brain MRI were created using a different GAN called DCGAN [20]. There was an association between experience with the data and the concordance between respondents (p=0.000). Between the first and second tests, the number of false positives decreased as expected, however, the number of false negatives increased, where 47% of all synthetic images were incorrectly reported as true, with 40% of false negatives in the expert group. These results further support that the synthetic prostate MRIs have realistic quality, and that the process for selection of higher quality synthetic prostate MRIs can be automated.

**Figure 3:**
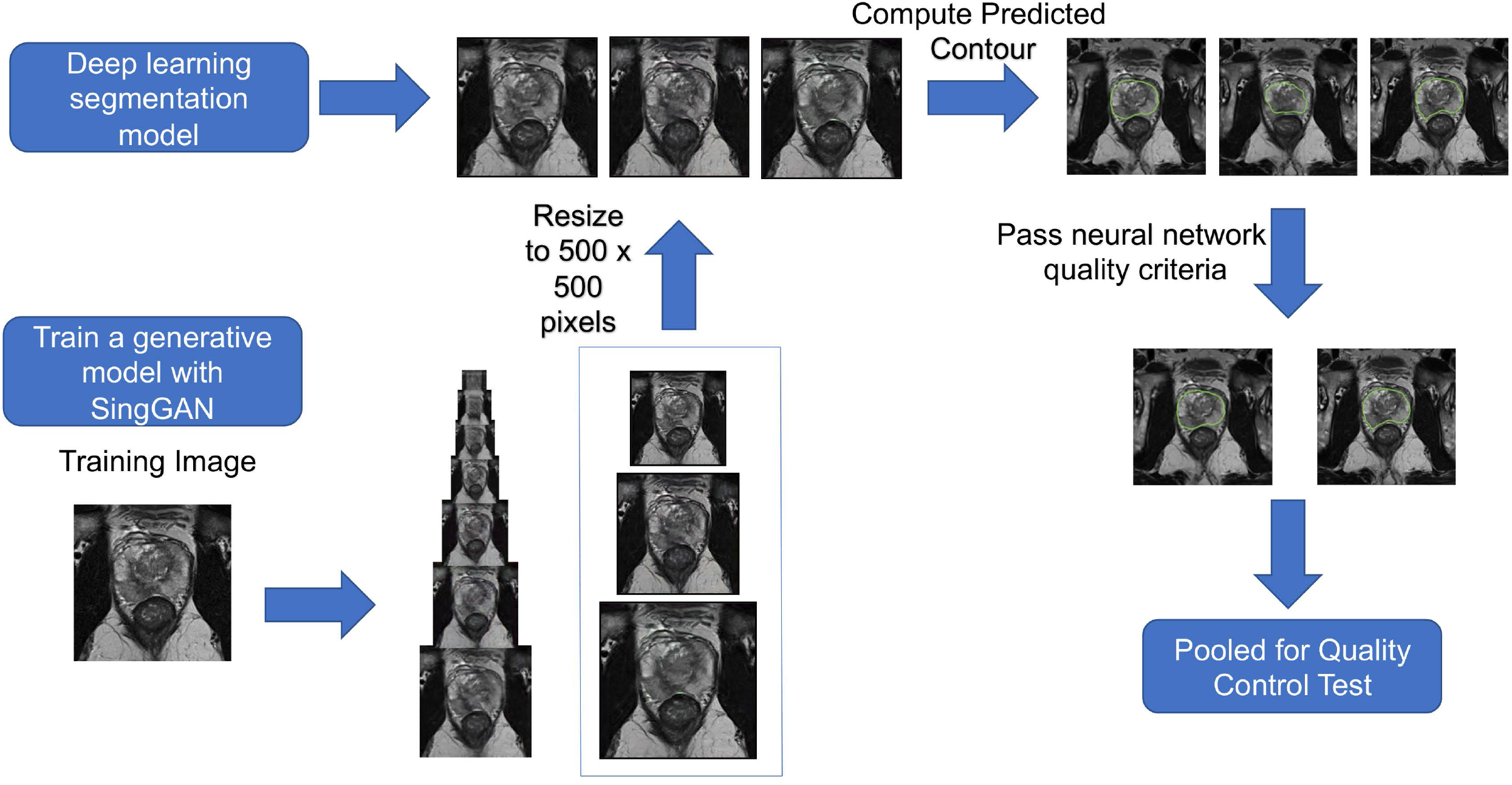
Workflow of procedure for second human quality control assessment. Three synthetic samples outputted by the last three generative models from SinGAN are resized to 500 × 500 then have a predicted segment of the prostate using a pre-trained segmentation neural network. Synthetic images that have a segmentation of the contour that pass our defined criteria were pooled and randomly selected for our human visual assessment.

### Radiologist Quality Control Check

To further assess the quality of the synthetic images, we enlisted the help of a board-certified radiologist to grade our prostate MRIs based on quality. Thirty images (20 synthetic, 10 conventional) that passed the criteria of the deep learning segmentation pipeline were randomly selected and given to the radiologist, who graded the images on a 1 to 10 scale based on the quality of being able to read and make a report (10 being best). The radiologist was NOT informed that the image set contained synthetic images. The mean grading was 6.2 for synthetic images and 5.5 for conventional images. A student t-test fails to reject the null hypothesis that the mean quality of synthetic and conventional images is equal (p=0.4839). The minimum, median, and maximum for the synthetic images and the conventional images are (1, 6, 10) and (1, 5.5, 10) respectively. These results suggest that the synthetic images are of equal quality when blindly given to a radiologist.

## Discussion

Our studies show that current machine learning technologies such as Generative Adversarial Networks can be used to generate synthetic prostate cancer MRIs that mimic conventional data. Other machine learning applications, such as deep learning semantic segmentation, can be used to automate the process of filtering out higher quality synthetic images. Our study demonstrated that participants did not have better performance on the quality control assessment when synthetic images were manually selected, and when the images were randomly pooled after passing the segmentation pipeline. This demonstrates the successfulness of the pipeline, because the human selection of the best synthetic images in the first visual assessment had no difference in scores from AI segmented synthetic images in the second assessment. Also, the synthetic images that passed the deep segmentation pipeline showed no difference compared to conventional images when graded based on quality by a board-certified radiologist. Our results show promise for future studies that involve synthetic imaging.

The filtering process for high quality synthetic images could be improved by using more training data in the segmentation neural network. Other studies involving automatic segmentation of the prostate with larger training databases show success in segmenting 2D and 3D prostate MRIs ^25^, as well as MRIs coming from multiple sources ^21^, such as our dataset. With only three slices each from 45 patients, the training data was relatively small and repetitive, which caused some visually realistic synthetic images not to pass the criteria. Furthermore, the criteria that the predicted prostate boundary had to be unbroken, disqualified many otherwise realistic looking synthetic MRIs, where other studies that included larger training databases also have predicted segmentations not one single boundary, but occasionally broken into pieces ^26^. Since deep learning segmentation is not completely accurate, quality scores can be given prediction boundaries that may be a better criterion in the future than the criteria used in this preliminary study ^26^. A segmentation model that considers other organs of the MRI such as bladder, urethra, gluteus maximus, rectum, or femur can also be implemented further select prostates with no distortion, although it will make the model highly selective and prone to overfitting. In turn, compressing the image more around the prostate can help reduce the amount of area the model needs to consider, and the amount of potential distortion, leading to enhanced segmentation. However, even with a very basic segmentation model, the model being able to segment the prostate proved an effective method in selecting high quality synthetic images. This is shown in Table 1, where 47% of synthetic images were mistaken for conventional, and 40% by experts. These results show promise of refinement of the model in future studies, where more accurate detection of prostate segmentation and or surrounding organs will enhance filtering of high-quality synthetic images.

In our study, the vendor used for the MRI and normalization programs were not stressed because we intended for participants to be presented with a wide variety of all types of MRI modalities that they may encounter. Furthermore, different versions of prostate MRI can be used such as dynamic contrast enhanced (DCE) perfusion imaging, where apparent diffusion coefficients (ADC) are commonly taken together ^27^, on top of the T2 diffusion weighted imaging used in this study, or PET scans, commonly used in other studies. The premise of this study can be extrapolated to these different image modalities, which may yield better results due to the decreased complexity of ADC images compared to T2W. Furthermore, the of the use of GAN in medical imaging studies is expanding to include 3-dimensional capabilities ^28^, where our study was limited to 2-dimensional slices of prostate MRI. Furthermore, a survey with more scientists and radiologists would increase the power of these results. There was no statistical difference in the rating of quality of synthetic and conventional MRI by the board-certified radiologist, but we stressed the promising result of the prostate cancer researcher’s due to their specificity with the prostate cancer MRI in particular. A different questioning method could be used similar to the quality of read recording we asked the board certified radiologist, or a Likert scale could be used as a scoring approach rather than binary responses ^29^. Additionally, our study had participants study images for 5 seconds, a longer assessment time per image should be tested to see if accuracy increases significantly with more consideration. Lastly, the 60 images used in the quality control tests contain only a small proportion of the total generated synthetic images (34 out of 253), so participants did not see the full array of synthetic images that passed the neural network criterion.

Lesion classification and tumor segmentation are also newfound capabilities of AI, where models such as random forest, naïve bayes, linear support vector machine (SVM), and CNNs have been able to show comparable Gleason grading accuracy to pathologists ^30^. These results support SinGAN’s unsupervised model using one training image, reducing tumor heterogeneity compared to models that combine multiple samples. Online databases such as those of the prostateX ^15^ challenge combine multiple views and modalities such as the ADC image, Sagittal, Coronal T2W images, on-top of the T2W images used in this study. They also include lesion data with all levels of Gleason grading that can be potentially classified using the above solutions in synthetic images. Future machine learning studies will be conducted to test if the high quality synthetic images can be integrated, and complement conventional images in classification, segmentation, and lesion detection.

In summary, we have demonstrated that synthetic prostate cancer MRI can be generated with high quality, mimicking conventional images. This process can be scaled to generate millions of unique synthetic samples. Other machine learning approaches such as the deep learning segmentation model can be used to remove synthetic images with high levels of distortion, which is one suggestion of the similar brain MRI study ^14^. This preliminary study suggests that synthetic prostate MR images may in the future be used in more complex imaging studies with clinical application.

## ACKNOWLEDGMENTS

We thank all the mentors, collaborators (Dr. Dipen J Parekh, Dr. Peter Clark, Dr. Joshua M Hare) and participants (Sameer Deshmukh, Yasamine Mirzabeigi, Brian Ledisma, Yaima Valdes, Ahmed Noman) for their insights, suggestions, and support during this study.

## Funding

This work was supported by American Urological Association Research Scholar Award for HA.

## Author contributions

HA, DVB, IX designed research; IX, DVB performed research; HA, IX, DVB, RS, SP, SG, AH analyzed data; and IX, HA, DVB wrote the manuscript.

## Competing interests

The authors declare no competing interests.

## Data and materials availability

All data, code, and materials used in the study are available to any researcher for purposes of reproducing or extending the analysis. Materials transfer agreements (MTAs) will be required to request the access.

## Notes

### Competing Interest Statement

The authors have declared no competing interest.

